# Phosphoantigens are Molecular Glues that Promote Butyrophilin 3A1/2A1 Association Leading to Vγ9Vδ2 T Cell Activation

**DOI:** 10.1101/2022.01.02.474068

**Authors:** Linjie Yuan, Xianqiang Ma, Yunyun Yang, Xin Li, Weiwei Ma, Haoyu Yang, Jian-Wen Huang, Jing Xue, Simin Yi, Mengting Zhang, Ningning Cai, Qingyang Ding, Liping Li, Jianxin Duan, Satish Malwal, Chun-Chi Chen, Eric Oldfield, Rey-Ting Guo, Yonghui Zhang

## Abstract

Tumor cells and pathogen-infected cells are presented to human γδ T cells based on “inside-out” signaling in which metabolites called phosphoantigens (pAgs) inside target cells are recognized by the intracellular domain of a butyrophilin protein (BTN3A1), leading to an extracellular conformational change. Here, we report that pAgs function as molecular “glues” that initiate a heteromeric association between the intracellular domains of BTN3A1 and the structurally similar BTN2A1. Working with both exogenous and endogenous pAgs, we used x-ray crystallography, mutational studies, cellular assays, synthetic probe as well as molecular dynamics investigations to determine how pAgs glue intracellular BTN3A1 and BTN2A1 together for the “inside-out” signaling that triggers γδ T cell activation. This γδ T cell-specific mode of antigen sensing creates opportunities for the development of alternative immunotherapies against cancer and infectious diseases that do not involve αβ T cells.

**One Sentence Summary:** The responses of gamma-delta T cells to cancer cells or pathogens are initiated via the intracellular association of heteromeric butyrophilins that are glued together by isoprenoid metabolites.

---

T cells recognize foreign molecules (antigens, Ags) via cell surface receptors (T cell receptors, TCRs) and provide immunity against tumor cells and pathogens. There are two families of T cells: αβ T cells and γδ T cells, defined by their TCRs. Knowledge about how αβ T cells recognize their antigens has expanded greatly over the last 40 years, including the realization that peptides are presented through major histocompatibility complex molecules (MHCs); that lipids are presented by CD1d to CD4, CD8, and natural killer T cells (NKTs); and that vitamin B metabolites are presented through MR1 to mucosal-associated invariant T cells (MAITs) (*1*). The common feature of Ag recognition by αβ T cells is that all antigens bind in a groove formed by an extracellular region of MHC-(like) antigen presenting molecules, enabling direct interactions with αβ TCRs.

In contrast to αβ T cells, the molecular basis for Ag recognition by γδ T cells is more enigmatic (*2*). Vγ9Vδ2 T cells, a major subtype of human circulating γδ T cells, respond to multiple cancers and infectious diseases in a MHC-independent manner: they are specifically activated by small, non-peptidic diphosphate metabolites called phosphoantigens (pAgs). Initial works in the mid-1990s showed that two small molecules, isopentenyl diphosphate (IPP) and dimethylallyl diphosphate (DMAPP), can weakly activate Vγ9Vδ2 T cells (*3*). Both of these “endogenous pAgs” (originating from inside the cell) are biosynthetic intermediates in the mevalonate pathway (*4*), and both are known to accumulate intracellularly during tumorigenesis (*5*). Later works identified a hydroxy-analog of DMAPP, (*E*)-1-hydroxy-2-methyl-but-2-enyl 4-diphosphate (HMBPP) as an exceedingly strong pAg of biological origin, with an EC_50_ for Vγ9Vδ2 T cell activation in the picomolar range (*6*). HMBPP is an “exogenous pAg” (originating from outside the cell) produced by gram-negative bacteria and by apicomplexan parasites via the methylerythritol 4-phosphate (MEP) pathway (*7*).

PAg sensing in both cancer cells and pathogen-infected cells is known to require a surface protein butyrophilin 3A1 (BTN3A1) (*8–12*), which has both intracellular and extracellular domains. Although the pAg binding site was initially thought to be located in the extracellular domain (*13*), this was later revised (*14–16*), and there is now crystallographic evidence that HMBPP binds to BTN3A1’s intracellular B30.2 domain (*14, 17*). This intracellular binding triggers an extracellular conformational change that allows target cells to be recognized by γδ T cells: “inside-out” signaling (*17*). Although now supported by multiple lines of evidence, this “inside-out” signaling mechanism with butyrophilins has only been previously demonstrated with integrin family proteins (*18*).

Structural and computational studies have indicated that a BTN3A1 monomer alone is unlikely to trigger the pAg-induced extracellular conformational change (*17*). It has therefore been assumed that either BTN3A1 dimers (*17, 19*) or complexes with one (or more) accessory protein(s) (*15, 16, 20–22*) are responsible for translating the intracellular pAg-recognition events to the extracellular changes which mark these cells for γδ T cell-mediated killing. We pursued the idea of contributions from additional protein(s) by carrying out a genome-wide CRISPR-Cas9 knockout screen using the cultured pancreatic cancer line MIA PaCa-2. By assaying the susceptibility of MIA PaCa-2 cells to Vγ9Vδ2 T cell mediated cell killing in the presence of the exogenous pAg HMBPP, we ultimately found that the loss-of-function for four butyrophilin family proteins protected MIA PaCa-2 cells from γδ T cell killing: BTN3A1, BTN3A2, BTN3A3, and BTN2A1 (fig. S1A). Both BTN3A2 and BTN3A3 belong to butyrophilin 3A family, and have been reported to function in immobilizing BTN3A1 (“3A1”) at the cell surface (*22*), thereby contributing to HMBPP-mediated Vγ9Vδ2 T cell killing. Of interest here, we also found that loss-of-function of BTN2A1 (BTN2A1^WT^ *versus* BTN2A1^-/-^) protected MIA PaCa-2 cells from HMBPP-mediated γδ T cell killing, the EC_50_ increasing from 6.5 nM to 36.0 µM (fig. S1B). The questions then arise: how might BTN2A1 function?

BTN2A1 (“2A1”) is predicted to comprise two extracellular domains (IgV and IgC), a single-pass transmembrane domain (TM), a proximal coiled-coil domain, and an intracellular B30.2 domain, followed by a C-terminal tail. It is known to interact with the C-type lectin receptor (CD209) in a glycosylation-dependent manner (*23*). The locus encoding 2A1 is on chromosome 6, which is consistent with a previous finding that Vγ9Vδ2 TCR-activation by pAgs requires 3A1 as well as additional genes on human chromosome 6 (*24*). Notably, two recent screening studies, one using CRSPR-Cas9 (*25*) and one using radiation hybrid screening (*26*), reported that 2A1 exerts an essential function in γδ T cell activation, mainly by using its extracellular IgV domain to directly interact with Vγ9^+^ TCR.

2A1 shares ∼50% sequence identity with 3A1’s intracellular B30.2 domain (fig. S1C), but isothermal titration calorimetry (ITC) showed that the recombinant 2A1 B30.2 domain did not bind to HMBPP (fig. S1D). This is in sharp contrast to the 1.64 μM *K*_D_ value detected for the HMBPP-3A1 B30.2 interaction (fig. S1D). We therefore determined the crystal structure of 2A1 B30.2 at 1.56 Å resolution (Table S1, PDB code: 7EQ8). The apo-form adopts the characteristic B30.2 fold that comprises a β-sandwich formed by two sets of antiparallel β sheets (sheets A and B) for its SPRY domain, together with a long loop (LTGANGVTPEEGLTLHRVSLLE) in the C-terminal tail (Fig. 1A). Monomeric 2A1 B30.2 can form a “head to tail” homodimer with another molecule by symmetry (Fig. 1B), with the dimer interface burying an area of approximately 1923.6 Å^2^ (accounting for 16% of the total surface area of the monomer) (fig. S1E). The extended C-terminal loop in each one monomer form part of the dimer interface (Fig. 1B). A subsequent analysis based on size-exclusion chromatography with multi-angle light scattering (SEC-MALS) showed that 2A1 B30.2 forms a dimer with a molecular weight of 50.98 kDa (∼25 kDa for each monomer) in solution, and truncation of the 2A1 B30.2 C-terminal tail diminished dimer formation (fig. S1F).

**Fig. 1.**
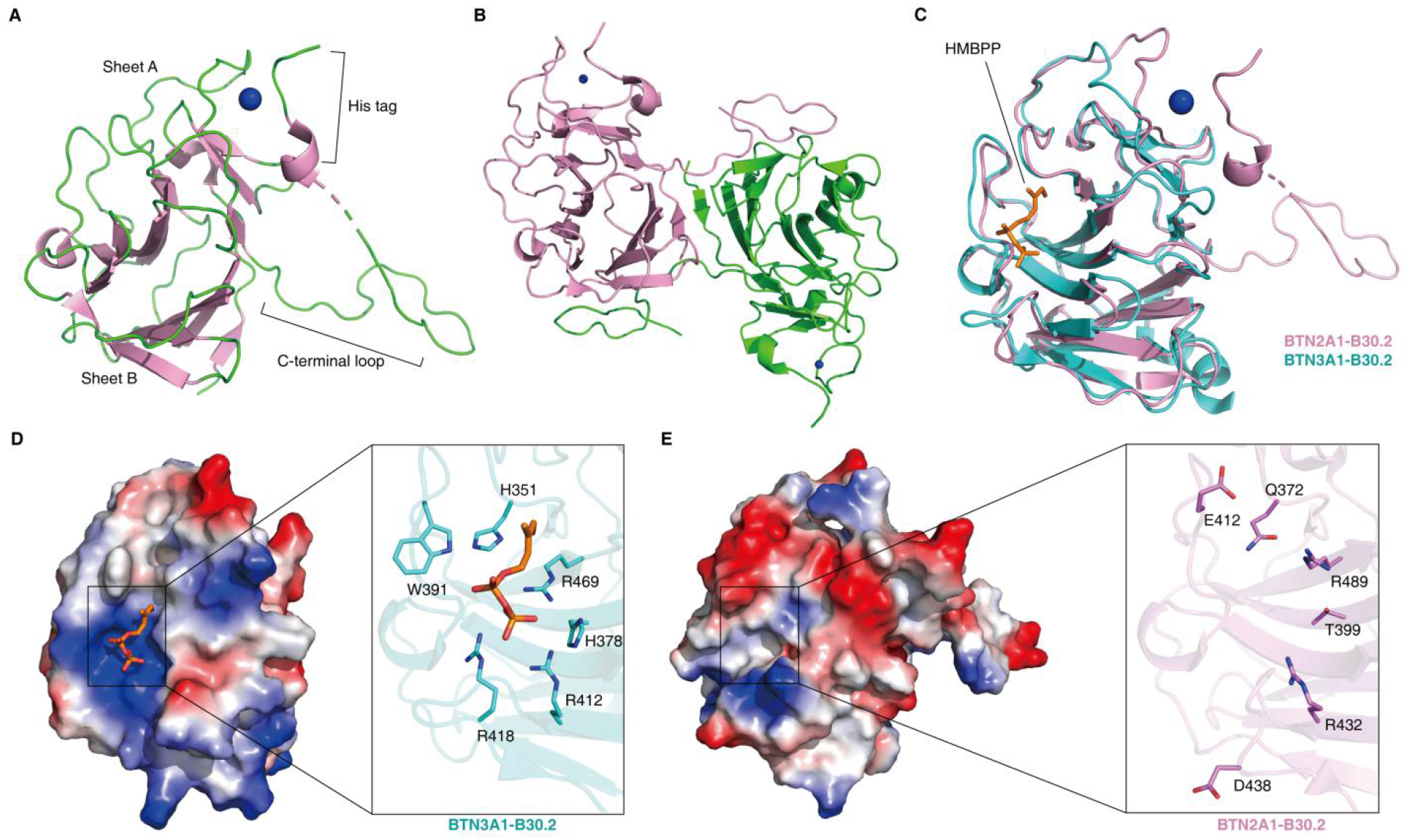
BTN2A1 is essential for HMBPP mediated Vγ9Vδ2 T cell activation but its B30.2 domain does not directly bind to HMBPP. **(A)** Cartoon model of apo-BTN2A1 B30.2 crystal structure (PDB code: 7EQ8). The two β sheets (sheets A and B) that comprise the core domain; the extended C-terminal loop; and the His tag are indicated. A zinc ion derived from His tag is displayed as a blue sphere. **(B)** The BTN2A1 B30.2 dimer. The two polypeptides chains are shown in pink and green. **(C)** Structure superimposition of BTN2A1 B30.2 (in pink) and HMBPP-bound BTN3A1 B30.2 (in cyan, PDB code: 5ZXK). The HMBPP observed in the BTN3A1 B30.2 structure is shown as a stick model. **(D)** Electrostatic surface of BTN3A1 B30.2 (left) and HMBPP-interacting residues (right). The highly cationic region is shown in blue. **(E)** Electrostatic surface (left) and residues (right) in BTN2A1 B30.2 structure corresponding to BTN3A1 B30.2 are illustrated.

The overall structure of the 2A1 B30.2 domain is very similar to that of the 3A1 B30.2 domain (PDB code: 5ZXK), with a Cα root-mean-square deviation (rmsd) of ∼0.67 Å (Fig. 1C). A major difference between the 2A1 and 3A1 B30.2 structures is that the 3A1 B30.2 domain has an HMBPP-binding basic pocket on the surface (*17*) (Fig. 1D), while no such basic pocket is evident in the corresponding region in the 2A1 B30.2 structure (Fig. 1E). The absence of the basic residues (three arginine and two histidine residues) that form the HMBPP-binding pocket in 3A1 obviously offers an explanation for the inability of 2A1’s B30.2 to bind HMBPP. The basic pocket and surrounding loops that bind HMBPP in 3A1 are positioned in its intracellular B30.2/PRYSPRY region (*17*) and it is of interest that the homologous regions in other B30.2-containing proteins have been shown to mediate protein-protein interactions (*27, 28*). Thus, although 2A1 B30.2 lacks the capacity to bind HMBPP by itself, it may undergo interactions with 3A1 B30.2.

We next used SEC-MALS to test whether 2A1 B30.2 can interact with 3A1 B30.2. In the absence of HMBPP, no interaction was detected. However, addition of HMBPP resulted in formation of 78.66 kDa complexes—likely comprising three monomeric B30.2 subunits of ∼25 kDa each (Fig. 2A). In addition, the ITC results showed that 2A1 B30.2 does not associate with 3A1 B30.2 in the absence of HMBPP. In the presence of HMBPP, 2A1 B30.2 does associate with 3A1 B30.2 with a *K*_D_ = 611 nM (Fig. 2B). These results suggest how HMBPP induces γδ T cell activation: HMBPP functions as a molecular glue that initiates an association between the B30.2 domains of 3A1 and 2A1. If so, this would help explain the requirement of 2A1 for γδ T cell activation (fig. S1, A and B).

**Fig. 2.**
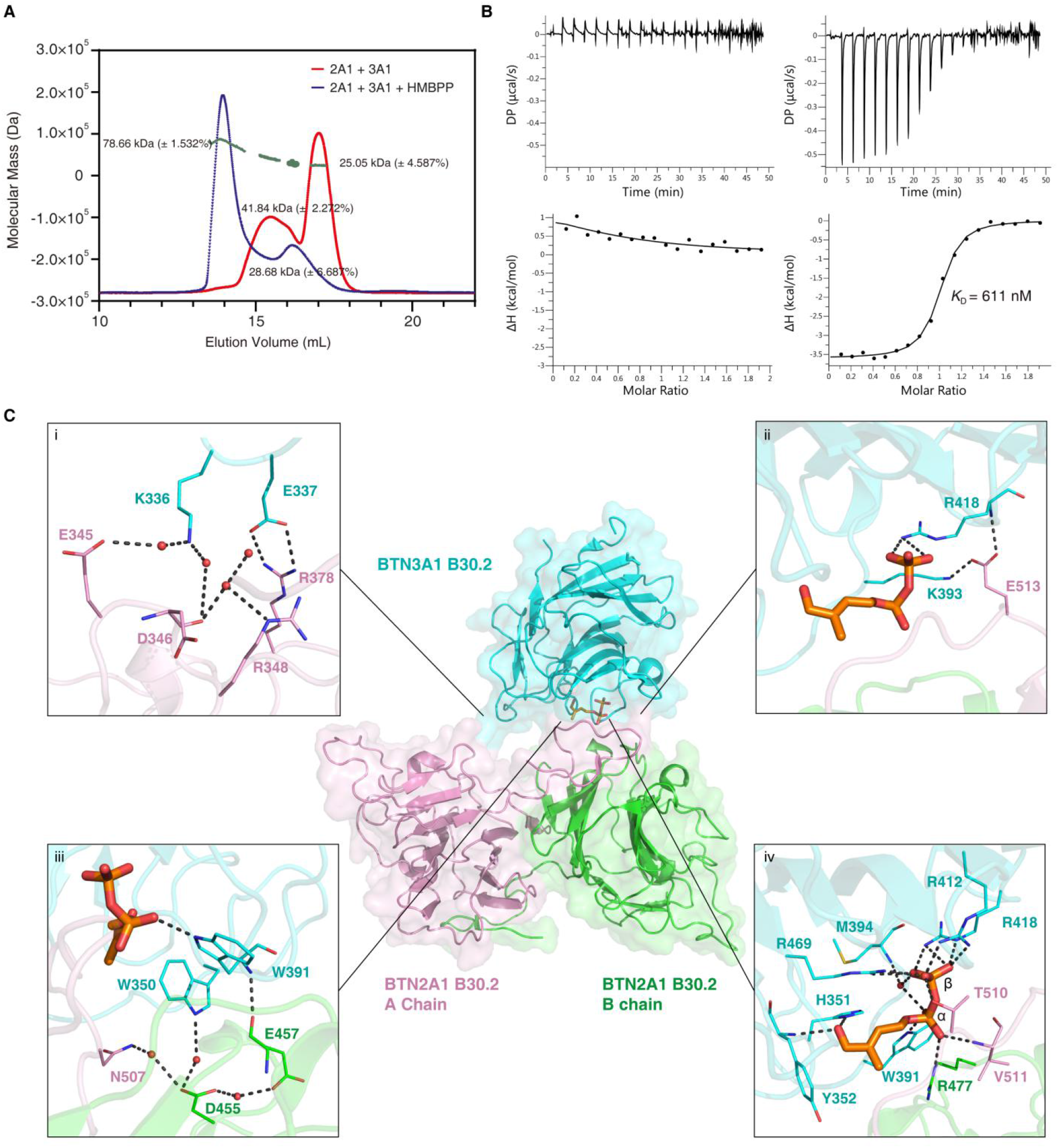
HMBPP acts as a molecular glue to promote the association of the BTN2A1 B30.2 and BTN3A1 B30.2 domains. **(A)** SEC-MALS analysis of 2A1 B30.2-3A1 B30.2 complexes with (in blue) or without (in red) HMBPP. **(B)** ITC results for 2A1 B30.2 binding to the 3A1 B30.2 domain with (right) or without (left) HMBPP. **(C)** Complex structure of 3A1 B30.2-HMBPP-2A1 B30.2 (PDB code: 7EQQ). Center: Cartoon representation of 3A1 B30.2-HMBPP (cyan) in complex with 2A1 B30.2 homodimer (pink and green for the A and B chains, respectively). HMBPP is shown as a stick model. Framed panels are close-up views of the interactions between **(i-ii)** 2A1 B30.2 and 3A1 B30.2 domains; **(iii)** W350/W391 (3A1 B30.2) and 2A1 B30.2 domain; and **(iv)** HMBPP and the 2A1-3A1 B30.2 domains. Protein residues and HMBPP are shown as lines and sticks, respectively. Waters are shown as small red spheres. Dashed lines indicate distance within 3.5 Å.

We thus next initiated a systematic effort to crystallize a complex between 3A1 B30.2, HMBPP and 2A1 B30.2, and ultimately solved a complex structure at 2.18 Å resolution (Table S1, PDB code: 7EQQ). There are two 2A1 B30.2 molecules and two HMBPP-bound 3A1 B30.2 molecules in one asymmetric unit. The HMBPP binding mode to 3A1 in the complex is similar to what was observed in our previously reported HMBPP-3A1 B30.2 complex crystal structure (PDB code: 5ZXK) (*17*) (fig. S2A). The two 2A1 B30.2 polypeptides constitute a homodimer basically as found in the apo-form 2A1 B30.2 structure. One 3A1 B30.2 molecule is located near to the 2A1 B30.2 dimer, with its HMBPP-binding pocket facing towards the C-terminal loop of the 2A1 B30.2 A chain and there are several polar interactions at the 3A1 and 2A1 interfaces. These include a salt bridge between E337 (3A1) and R378 (A chain, 2A1) (Fig. 2C, i), and a salt bridge between K393 (3A1) and E513 (A chain, 2A1) (Fig. 2C, ii). In addition, there is also a hydrogen bond between R418 (3A1) and E513 (A chain, 2A1) (Fig. 2C, ii).

Previous MD simulations indicated functional contributions from 3A1’s W350 and W391 residues in the transmission of the “inside-out” signal induced by HMBPP binding (*17*), so it is of interest to note that the 3A1 B30.2-HMBPP-2A1 B30.2 structure also shows interactions among these C-terminal tail residues (Fig. 2C). Specifically, W391 (in 3A1) forms a hydrogen bond (at 2.9 Å) with E457 (2A1, B chain), while W350 (3A1) forms a water-mediated hydrogen bond (∼3 Å) with D455 (2A1, B chain) (Fig. 2C, iii). It should be noted that the above findings (Fig. 2C, i-iii) do not unambiguously establish that these interactions must be induced by HMBPP binding.

The most striking finding from the 3A1 B30.2-HMBPP-2A1 B30.2 structure relates to the HMBPP molecule: its Pα phosphate and Pβ phosphate each undergo strong interactions with 2A1 (Fig. 2C, iv). The Pα phosphate forms two interactions (one hydrogen bond at 2.7 Å and one salt bridge at 3.9 Å) with R477 (2A1, B chain), and one hydrogen bond with V511 (2A1, A chain, at 2.9 Å). The Pβ phosphate has one hydrogen bond between its oxygen and T510 (2A1, A chain, at 2.7 Å).

These structural findings support the hypothesis that HMBPP functions as a molecular glue to initiate an intracellular 2A1 and 3A1 association. To test whether HMBPP-mediated 2A1/3A1 association results in Vγ9Vδ2 T cell activation, we next mutated 2A1 R477, T510 and V511, each of which directly interacts with HMBPP. Unlike wild type 2A1, none of these 2A1 mutants associated with 3A1 or caused Vγ9Vδ2 T cell activation (fig. S2, B and C), supporting the hypothesis that it is the “glue” role of HMBPP which leads to γδ T cell activation.

We also used mutagenesis to confirm functional contributions to Vγ9Vδ2 T cell activation. Briefly, in previous work we found that W350 and W391 in 3A1 were required for HMBPP-mediated inside-out signaling (*17*), but the precise nature of their contribution(s) was not clear. Since the 3A1 B30.2-HMBPP-2A1 B30.2 structure revealed interactions between the 3A1 W350/W391 residues and 2A1’s D455, E457 residues (Fig. 2C, iii), we generated a 2A1 double mutant D455G/E457R to test the importance of this interation. This mutant lost its ability to mediate HMBPP-triggered Vγ9Vδ2 T cell activation (fig. S2C). Also, given that the large majority of 2A1’s interactions with 3A1 involve its C-terminal tail, we truncated this tail, finding that it abrogated the HMBPP-induced 3A1-2A1 association and abolished the HMBPP-mediated susceptibility of MIA PaCa-2 cells to Vγ9Vδ2 T cell killing (fig. S2, B and C). Collectively, the structural and mutational results support the hypothesis that the exogenous pAg HMBPP functions as a natural molecular glue which mediates γδ T cell activation by initiating the association of the B30.2 domains of 2A1 and 3A1 inside target cells.

One obvious question extending from this pAg molecular glue hypothesis relates to the capacity of endogenous pAgs (DMAPP and IPP) to initiate association of 3A1 and 2A1. The FDA approved bisphosphonate drug zoledronate activates γδ T cells via accumulation of DMAPP and IPP (*29*). The Uldrich group reported that 2A1 was also required for zoledronate-induced γδ T cell activation (*25*), indicating its essentiality for endogenous pAgs, which we confirmed in the present study (fig. S3A). ITC analysis indicated that 3A1 B30.2 alone has a poor binding affinity (120 μM) for DMAPP (Fig. 3A) (vs. 1.64 μM for HMBPP), perhaps explaining our failure to obtain a crystal structure of a DMAPP-3A1 B30.2 complex. However, DMAPP promoted the association of 3A1 with 2A1 B30.2 (*K*_D_ = 12.3 μM) (Fig. 3A). So even though its binding is substantially weaker than HMBPP (12.3 µM vs. 0.6 µM), these results show that the endogenous pAg DMAPP can also function as a molecular glue to promote the 2A1-3A1 B30.2 association. In contrast, IPP did not promote the 2A1-3A1 B30.2 association (Fig. 3A), suggesting that DMAPP may be the more potent phosphoantigen (*3*).

**Fig. 3.**
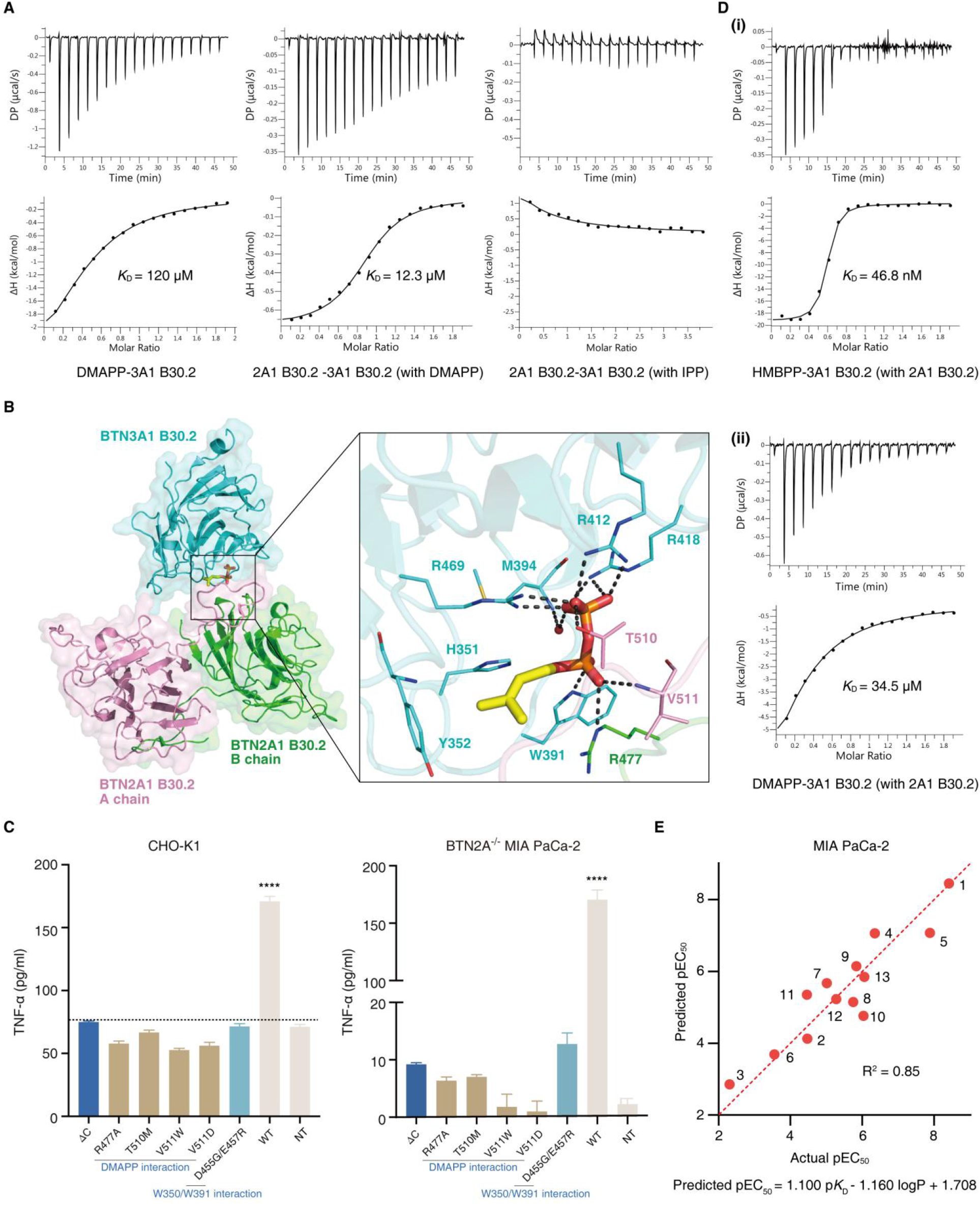
The endogenous pAg DMAPP and synthetic pAgs also function as molecular glues. **(A)** ITC results for DMAPP binding to the 3A1 B30.2 domain (left) and the 2A1 B30.2 domain binding to the 3A1 B30.2 domain with DMAPP (middle) or IPP (right). **(B)** Cartoon model of the 3A1 B30.2-DMAPP-2A1 B30.2 complex. One 3A1 (cyan) and two 2A1 B30.2 domains (pink and green for the A and B chains, respectively) are aligned as in the 3A1 B30.2-HMBPP-2A1 B30.2 complex (Fig. 2C). DMAPP is shown as a stick model. The close-up view illustrates the interaction networks formed by DMAPP and the three polypeptide chains. Protein residues are shown as lines, a water is shown as a red sphere. **(C)** TNF-α release by Vγ9Vδ2 T cells in response to zoledronate (Zol, 10 µM) stimulation of 3A1^+^CD80^+^ CHO-K1 cells (left, *n* = 5) or BTN2A^-/-^ MIA PaCa-2 (2A1/2A2 KO) cells (right, *n* = 5) transfected with the plasmids for the indicated 2A1 mutants. Residues in 2A1 B30.2 directly interacting with DMAPP are in brown. Residues in 2A1 B30.2 directly interacting with W350/W391 in 3A1 B30.2 are in cyan. The horizontal dotted line indicates TNF-α release of NT, which was a blank plasmid used as a control for comparison. Data were analyzed by using two-tailed unpaired *t* tests. \*\*\*\**P* < 0.0001. All error bars denote the SEM. **(D)** ITC results for HMBPP (i) and DMAPP (ii) binding to the pre-conditioned 2A1-3A1 B30.2 complex. **(E)** Correlation between experimental and predicted pEC_50_ values (pEC_50_= - log_10_EC_50_ [M]) for phosphoantigen-mediated Vγ9Vδ2 T cell killing of MIA PaCa-2 cells, obtained by using experimental *K*_D_ (2A1/3A1 assay; Table S2) and computed logP (Table S2) values for a library of HMBPP analogs (fig. S3C).

Despite our failure to crystallize a DMAPP-3A1 B30.2 complex, we successfully determined a 3A1 B30.2-DMAPP-2A1 B30.2 complex structure at 2.29 Å resolution (Table S1, PDB code: 7EQY). The complex structure closely resembles the 3A1 B30.2-HMBPP-2A1 B30.2 complex (Fig. 3B and Fig. 2C) but in contrast to HMBPP, DMAPP does not form two hydrogen bonds with 3A1’s H351 and Y352 residues (Fig. 3B), due of course to the absence of a 1-OH group. The lack of these two hydrogen bonds means that DMAPP binds more weakly to 3A1 which then results in weaker binding to 2A1. In the presence of DMAPP, we see that the intramolecular interactions between 2A1 and 3A1 B30.2 are virtually the same with HMBPP (fig. S3B). Moreover, the same 2A1 mutations which disrupted Vγ9Vδ2 T cell activation in the presence of HMBPP also impaired Vγ9Vδ2 T cell activation in experiments using zoledronate-treated CHO-K1 cells or BTN2A^-/-^ MIA PaCa-2 cells (Fig. 3C). All of these results support the idea that DMAPP functions as an endogenous molecular glue in tumor cells that leads to γδ T cell activation.

This molecular glue hypothesis opens new opportunities to investigate the puzzling fact in γδ T cell biology that the cellular activities of pAgs (as low as pM levels) do not correlate with their 3A1 B30.2 binding affinities (μM range). That is, guided by the observation that pAgs function as molecular glues at the interface formed by 3A1 and 2A1 B30.2, we are now better positioned to design experiments to study the relationship between pAg cellular activities and binding affinities. We thus next conducted a series of binding assays with natural pAgs as well as with a series of synthetic HMBPP analogs to help to clarify this aspect of inside-out Vγ9Vδ2 T cell activation. We first incubated 2A1 B30.2 and 3A1 B30.2 proteins (at a 2:1 2A1/3A1 ratio) and used ITC to measure HMBPP binding affinity for this “pre-conditioned complex”. The *K*_D_ value for HMBPP went from 1.64 µM for 3A1 alone to 46.8 nM for the 2A1-3A1 B30.2 complex (fig. S1D and Fig. 3D, i), a value much closer to the observed picomolar activity of HMBPP in Vγ9Vδ2 T cell activation. For DMAPP, the *K*_D_ values went from 120 μM to 34.5 μM (Fig. 3A and Fig. 3D, ii), consistent with the fact that DMAPP is a far weaker Vγ9Vδ2 T cell activator. We next investigated a library of HMBPP analogs (structures shown in fig. S3C), measuring their binding affinities to the pre-conditioned 2A1-3A1 B30.2 complex, as well as their activities for sensitizing MIA PaCa-2 cells to Vγ9Vδ2 T cell killing (Table S2). We found a very good correlation (R^2^ = 0.85) between the experimentally observed Vγ9Vδ2 T cell killing activities (pEC_50_ = -log_10_EC_50_[M]) when using the experimental *K*_D_ and computed logP values (Fig. 3E).

We next conducted molecular dynamics (MD) simulations to explore how these molecular-glue mediated intracellular associations are transmitted outwards to evoke extracellular signaling events. Using the two 3A1 B30.2-pAgs-2A1 B30.2 complex structures, with or without HMBPP or DMAPP, we found that plotting the Cα root-mean-square fluctuations (RMSF) for the trajectories identified a set of residues with relatively large fluctuations (defined as RMSF>1.5 Å, and shown in red in fig. S3D. Full trajectories are shown in fig. S3E). The simulations clearly indicate that the 2A1 B30.2 subunits remain rigid—either with or without pAgs—whereas 3A1 B30.2 is more flexible when no pAg is present (fig. S3E). Further, binding of pAgs to the trimeric complex is able to pass the binding fluctuations to the coiled-coil α-helical regions (juxtamembrane) known to link the B30.2 domains to the cell membrane (*30*) (fig. S3D).

Finally, we investigated the role of “the other side of the cell membrane” by examining structurally-related butyrophilins. Both mutagenesis and protein-region-engineering studies have revealed specific contributions of particular butyrophilin residues and regions to “inside-out” pAg presentation to γδ T cells. As an example, the B30.2 domain of BTN2A2 (“2A2”) shares 88.7% similarity with 2A1 B30.2 (fig. S4A); however, we found that 2A2 B30.2 does not associate with 3A1 B30.2, even in the presence of HMBPP (Fig. 4A). Systematic screening of diverse 2A2 mutant variants (with residues selected for mutagenesis based on their homology to residues in 2A1 B30.2 and 3A1 B30.2) revealed that mutants of 2A2 B30.2 bearing a M506T mutation or two mutations (W374R/M506T) were able to interact with 3A1 B30.2 (in the presence of HMBPP) with *K*_D_ values of 353 nM and 189 nM, respectively (Fig. 4A). M506 and W374 are analogous to 2A1 B30.2 residues T510 and R378, which we earlier found were in direct interactions with pAgs.

**Fig. 4.**
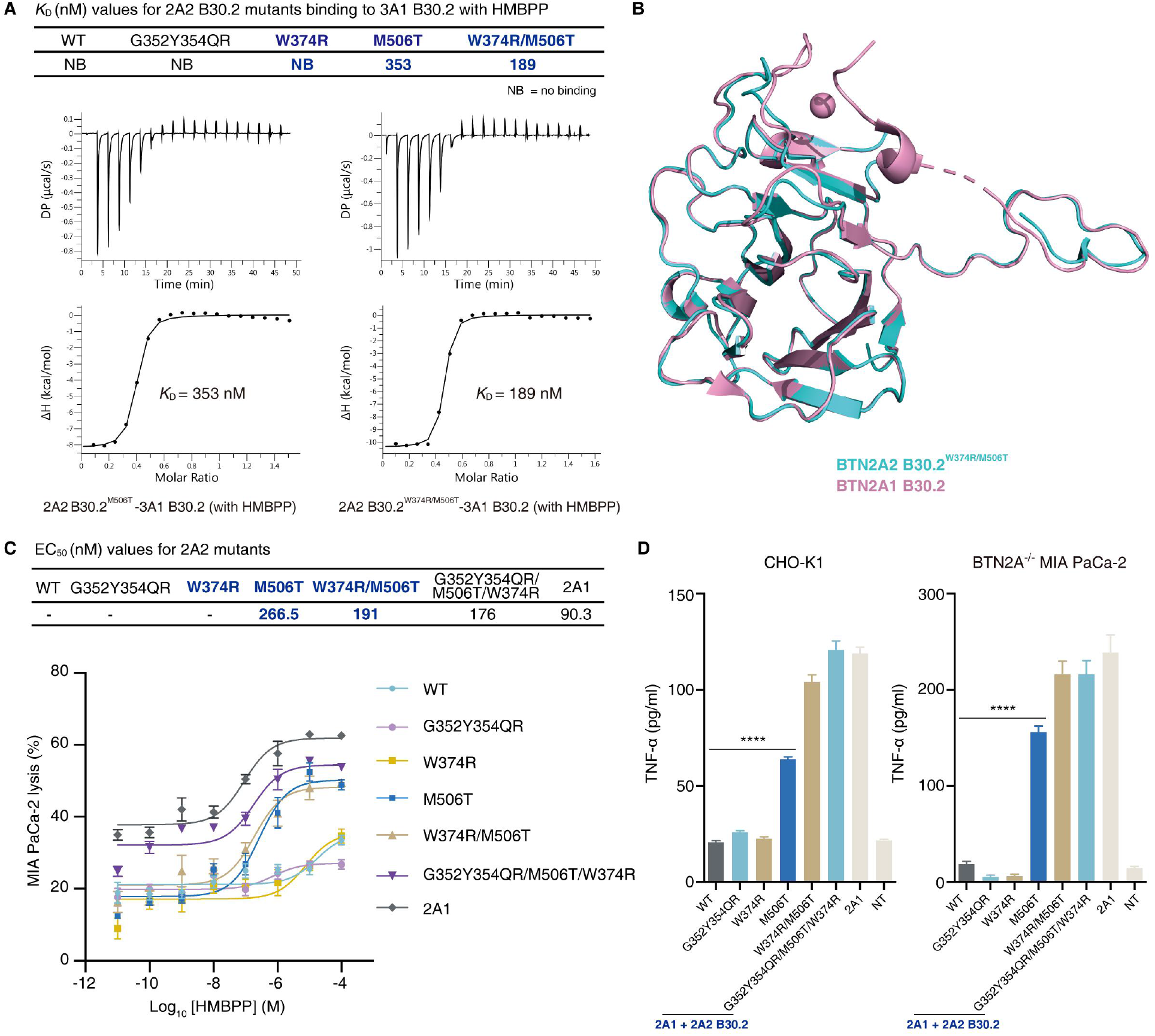
Both the intracellular and extracellular regions of BTN2A1 are required for “inside-out” signaling for γδ T cell activation. **(A)** Top: *K*_D_ values (nM) for WT, single and double 2A2 B30.2 mutants binding to 3A1 B30.2 with HMBPP. Bottom: W374 and M506 are the 2A2 residues that align to the 2A1 B30.2 residues which interact directly with 3A1 B30.2 or with HMBPP, as seen in the 3A1 B30.2-HMBPP-2A1 B30.2 complex structure (PDB code: 7EQQ). Note that 2A1 has an additional loop (C353F355HLGES) which is not present in 2A2 (G352Y354QR). Representative ITC calorimetry plots characterizing two 2A2 B30.2 mutants (M506T and W374R/M506T) binding to the 3A1 B30.2 domain in the presence of HMBPP. **(B)** Structural superimposition of the 2A2 B30.2 ^W374R/M506T^ domain (cyan, PDB code: 7ER4) and the 2A1 B30.2 domain (pink, PDB code: 7EQ8). **(C**) Top: Cytotoxicity (nM) of Vγ9Vδ2 T cells towards BTN2A^-/-^ MIA PaCa-2 (2A1/2A2 KO) cells transfected with plasmids for the indicated combinations of 2A1 + 2A2 B30.2 domain mutant variants treated with HMBPP (10 pM to 100 µM). Bottom: Dose-response curves for MIA PaCa-2 cell lysis by Vγ9Vδ2 T cells treated with HMBPP. **(D)** TNF-α release by Vγ9Vδ2 T cells in response to Zol (10 µM) stimulation of 3A1^+^CD80^+^ CHO-K1 cells (left, *n* = 6) or BTN2A^-/-^ MIA PaCa- 2 cells (right, *n* = 4), transfected with the plasmids for the indicated combinations of 2A1 + 2A2 B30.2 domain mutants (2A1 extracellular domain, 2A1 TM, 2A1 coiled-coil, and 2A2 B30.2 mutants). Data were analyzed by using two-tailed unpaired *t* tests. \*\*\*\**P* < 0.0001. All error bars denote the SEM.

We determined the crystal structure of the 2A2 B30.2^W374R/M506T^ variant at 2.12 Å resolution (Table S1, PDB code: 7ER4). Even though this structure was almost identical to that of 2A1 B30.2 (Fig. 4B), cellular assays showed that this 2A2 mutant variant was not able to induce Vγ9Vδ2 T cell activation (fig. S4, B and C). Moreover, protein engineering work with 2A2 showed the essential contributions of multiple 2A1 extracellular structures to γδ T cell activation: it was only when we replaced 2A2’s coiled-coil, TM and extracellular domains with their 2A1 counterparts that any Vγ9Vδ2 T cell activation was detected (Fig. 4, C to D and fig. S4, B to C). These results are consistent with the pioneering work from the Uldrich and Herrmann groups, which established a role for 2A1’s extracellular domain as a direct ligand for the γδ TCR (*25, 26*). Collectively, these results demonstrate that both the intracellular and extracellular domains of 2A1 are required for Vγ9Vδ2 T cell activation (fig. S4D).

In summary, the structural and cellular assay results from the present study establish that both exogenous and endogenous phosphoantigens act as molecular glues to promote the butyrophilin 3A1/2A1 association that initiates signaling within target cells, leading to Vγ9Vδ2 T cell activation. These basic insights can inform rational design of specific modulations for γδ T cell-based immunotherapies to treat cancers and infectious diseases. More broadly, consider that a B30.2 protein-binding domains are present in the intracellular tail of most butyrophilins, heteromeric interactions are often required for their immune functions (*22, 31, 32*), and multiple studies of structurally-related butyrophilins have reported diverse immunomodulatory activities (*31*), we therefore anticipate that our discovery of pAgs as molecular glues linking 3A1/2A1 inside target cells will inform investigations to elucidate the roles of other butyrophilins in immune modulations.

## Acknowledgments

We thank David Huang (University of Melbourne) for providing the whole-genome sgRNA library plasmid; Erin Adams (University of Chicago) for providing us with the BTN3A1 B30.2 domain plasmid; Dr. Shilong Fan (Tsinghua University) and the staff at the Shanghai Synchrotron Radiation Facility (SSRF) for assistance in data collection.

## Author contributions

Y.Z. initiated the project and designed experiments. L.Y., X.M., Y.Y., X.L., W.M., H.Y., J.-W.H., J.X., S.Y., M.Z., N.C., Q.D. and L.L. performed experiments. L.Y., H.Y. and Q.D. carried out CRISPR-Cas9 knock out screening. L.Y. carried out proteins cloning, expression and crystallizations of BTN2A1 and BTN3A1 B30.2. Y.Y., X.M., M.Z. and S.Y. carried out proteins cloning, expression of BTN2A1 and BTN2A2 B30.2 mutants and crystallizations of BTN2A2 B30.2 mutant. Y.Y., X.L, N.C., J.-W.H. and C.-C.C contributed to crystallization data analysis. L.Y., X.M., and Y.Y. carried out full length mutation plasmid cloning. L.Y. performed cell experiments. X.M. and L.L. synthesized all chemicals. X.M. and L.Y. carried out ITC experiments. X.M. and J.D. performed computational studies. X.M. and Y.Y. performed SEC-MALS experiments. W.M. performed chemicals activities tests. Y.Z. supervised the whole research and wrote the paper, with comtributions from R.-T.G., L.Y., Y.Y., S.M. and E.O. to experiments design and discussions.

## Data and materials availability

The crystal structures are deposited in the Protein Data Bank under accession code 7EQ8, 7EQQ, 7EQY and 7ER4 respectively. All data needed to evaluate the conclusions in the paper are present in the paper and Supplementary Materials.

## Materials and Methods

### Cell lines

MIA PaCa-2 cells, a human pancreatic cancer cell line, were purchased from ATCC. HEK293T cells, a human embryonic kidney cell line, were purchased from ATCC. MIA PaCa-2 cells and 293T cells were cultured in Dulbeco’s Modified Eagle’s medium (DMEM) (Gibco, cat. no. 11960-051) supplemented with 10% heat inactivated FBS (Biolnd, cat. no. 04-001-1), 1% penicillin/streptomycin (Beyotime, cat. no. C0222) at 37 ℃ and 5% CO_2_. CHO-K1 cells, a subclone from the parental CHO cell line initiated from a biopsy of an ovary of an adult Chinese hamster, were purchased from ATCC. CHO-K1 cells were cultured in Ham’s F12K (Kaighn’s) medium (HyClone, cat. no. SH30526.01) supplemented with 10% heat inactivated FBS, 1% penicillin/streptomycin at 37 ℃ and 5% CO_2_.

### Protein expression and purification

The BTN3A1 B30.2 domain plasmid was provided by Professor Erin Adams of the University of Chicago. BTN3A1 B30.2 was expressed and purified as described previously (*14, 17*). The cDNA encoding the BTN2A1 B30.2 domain was cloned into the pET28a vector with N-terminal or C-terminal 6 × His tags, and cDNA for the BTN2A2 B30.2 domain was also cloned into the pET28a vector with N-terminal 6 × His tags. Overexpression of the BTN2A1 B30.2 domain was induced in *E. coli* BL21 (DE3) cells by 1 mM isopropyl β-D-1-thiogalactopyranoside (IPTG) (Solarbio, cat. no. 367-93-1) at 18 ℃ for 24 h. Cells were collected by centrifuging, resuspended and lysed by sonication, and centrifuged at 20,000 g for 1 h. The supernatant was collected and incubated with Ni-NTA resin (GE Healthcare), and was then eluted with a buffer containing 20 mM HEPES pH 7.5, 150 mM NaCl, and 500 mM imidazole. The eluent was incubated overnight with TEV protease for hexahistidine tag cleavage, purified by Ni-NTA resin and further purified by DEAE Sepharose chromatography (GE Healthcare). Finally, the protein was dialyzed against storage buffer containing 20 mM HEPES pH 7.5, 150 mM NaCl. All BTN2A1 B30.2 domain mutants were generated with a standard PCR mutagenesis strategy, overexpressed, and purified in the same way as for the wild-type protein. The expression and purification of BTN2A2 B30.2 WT and mutants followed the same methods as for the BTN2A1 B30.2 domain.

### Crystallization, data collection, and structure determination

All crystals were obtained using the sitting-drop vapor diffusion method. Apo-BTN2A1 B30.2 domain crystals appeared in buffer containing 0.4 M potassium sodium tartrate. The crystals of BTN2A1-BTN3A1 B30.2 with HMBPP or DMAPP appeared in buffer containing 42% v/v polyethylene glycol 200 and 0.1 M HEPES pH 7.5. All data were collected at the Shanghai Synchrotron Radiation Facility (SSRF), beamlines BL17U1, BL18U1, and BL19U1, and data were processed using the HKL-2000 program. BTN2A2 B30.2^W374R/M506T^ domain crystals appeared in buffer containing 0.2 M sodium citrate and 20% w/v polyethylene glycol 3350. X-ray diffraction data were obtained at the in-house beamline BRUKER D8 VENTURE at Hubei University. Datasets were processed with PROTEUM3 (Bruker AXS GmbH). Data collection and structure refinement statistics are shown in Table S1. All structures were solved by molecular replacement using the BTN3A1 B30.2-HMBPP complex structure (PDB code: 5ZXK) as the search model, and were refined with Coot and Refmac (*33, 34*).

### Vγ9Vδ2 T cell isolation and expansion

All studies using Vγ9Vδ2 T cells were carried out in accordance with the recommendations of the Institutional Review Board of Tsinghua University with written informed consent from all subjects. All subjects gave written informed consent in accordance with the Declaration of Helsinki. The protocol was approved by the Institutional Review Board of Tsinghua University (Project no: 20170004 and 20170007). Peripheral blood mononuclear cells (PBMCs) were isolated from healthy donors by density gradient centrifugation using Ficoll-Hypaque density fluid (GE Healthcare, cat. no. 17144003). PBMCs were cultured at a density of 2 × 10^6^ cells/mL in RPMI 1640 (Gibco, cat. no. 27016-021) that was supplemented with 10% FBS, 1% penicillin-streptomycin, 150 U/mL human rIL-2 (PeproTech, cat. no. 96-200-02), 1% MEM non-essential amino acids (Gibco, cat. no. 11140050), 2 mM L-glutamine (Beyotime, cat. no. C0212), 50 mM β-mercaptoethanol (Amresco, cat. no. 0482), and 5 mM zoledronate (Energy Chemical, cat. no. A010559) at 37℃ and 5% CO_2_. Fresh medium containing human IL-2 was replaced every 3 days. Vγ9Vδ2 T cells were harvested at 11 days and stored for future use.

### Viral library production

The whole-genome sgRNA library plasmid (GeCKOv2, a gift from David Huang, University of Melbourne) plus packaging plasmids using FuGENE6 (Promega, cat. no. E2691) were transfected into low-passage HEK293T cells at 80% confluency in 15 cm tissue culture dishes. Viral supernatants were collected at 48-72 h post-transfection, filtered through a 0.45 µm filtration unit, and stored at -80 ℃ for future use. Virus for other plasmids (including cas9, BTN3A1 and CD80) were produced in a similar manner.

### Whole-genome CRISPR-Cas9 knockout screen

The CRISPR-Cas9 knockout screen was performed essentially as described (*35–37*). MIA PaCa-2 cells were infected with the lentiviral supernatant containing cas9 at approximately 80% confluence in the presence of polybrene, and cas9 positive MIA PaCa-2 cells were then sorted by FACS 48 h later. Sorted cells were infected with the lentiviral-packaged whole-genome sgRNA library to achieve 30% transduction efficiency, and transduced cells were selected with puromycin for an additional 10 days. MIA PaCa-2 cells (> 100 × library coverage) were pretreated with HMBPP (10 nM) for 4 h, and then co-incubated with Vγ9Vδ2 T cells at a T cell: cancer cell ratio = 10:1. After 72 h, Vγ9Vδ2 T cells were removed and remnant MIA PaCa-2 cells were re-expanded for 1-2 weeks. This process was repeated an additional 10 consecutive times. Finally, genome sequencing was conducted for cells at each step, and sgRNA counting was performed and analyzed as reported (*37, 38*). The raw count files and analysis script are available in database S1.

### Generation of MIA PaCa-2 BTN2A1/BTN2A2 knockout cells

BTN2A1 and BTN2A2 genes were disrupted in MIA PaCa-2 cells using CRISPR. The CRISPR sequencing targeting functional BTN2A genes were designed with the help of online tools (http://crispr.mit.edu) mentioned in Table S3 and sequences were cloned into a PX458-pSpCas9(BB)-2A-GFP-MCS vector. All plasmids were sequenced to confirm successful ligation. The plasmids were then transfected into MIA PaCa-2 cells using Lipofectamine 2000 reagent (Invitrogen, cat. no.11668-019). Single cell clones were sorted (GFP), and were then validated based on genomic sequencing.

### Cell killing assays

MIA PaCa-2 cells were infected with lentivirus bearing luciferase transgene. Luciferase-positive cells were plated at 5.0 × 10^3^ cells/well in 384 well plates one day prior to transfection. All plasmids were transfected into MIA PaCa-2 cells using Lipofectamine 2000 reagent. After 48 h, MIA PaCa-2 cells, which were pretreated with different concentrations of HMBPP for 4 h, were co-cultured with 1.0 × 10^4^ Vγ9Vδ2 T cells for 16 h. Stable-lite^TM^ luciferase (Vazyme, cat. no. DD1202-01) was added to each well and the luciferase signal was immediately measured by using a PerkinElmer EnVision.

Percent lysis was calculated by using the following equation: % specific lysis = (Maximum Luciferase - Experimental Luciferase) / Maximum Luciferase

### TNF-α secretion assays of Vγ9Vδ2 T cells co-cultured with BTN2A^-/-^ MIA PaCa-2 cells or CHO-K1 cells

BTN2A^-/-^ MIA PaCa-2 (2A1/2A2 KO) were plated at 1.0 × 10^4^ cells/well in 96 well plates 1 day prior to transfection. All plasmids were transfected into MIA PaCa-2 cells using Lipofectamine 2000 reagent, followed with treatment with zoledronate (10 µM) for 24 h. Then the medium was aspirated and cells were washed 4 times with PBS at room temperature. Vγ9Vδ2 T cells (1.0 × 10^5^ cells/well) were then added and cocultured with MIA PaCa-2 cells, and culture supernatants were harvested after 16 h and assayed for TNF-α levels using a TNF-α human uncoated ELISA kit (Invitrogen, cat. no. 85-88-7346). The assay of CHO-K1 cells (infected with lentivirus of 3A1 and CD80) used the same methods as MIA PaCa-2 cells (*25, 26*).

### Isothermal titration calorimetry (ITC)

ITC experiments were carried out using a MicroCal PEAQ-ITC instrument (GE Healthcare). We used an initial injection of 0.4 µL followed by 19 injections (2 µL each) at 150 s time intervals. ITC binding fits were calculated using Microcal Origin software. Samples were prepared using a buffer (20 mM HEPES pH 7.5, 150 mM NaCl).

For pAgs binding to the BTN3A1 B30.2 domain, the sample cell and injection syringe were filled with the buffer containing 100 µM protein and 1 mM HMBPP respectively, or 400 µM protein and 4 mM DMAPP (IPP) respectively.

For pAgs binding to the pre-conditioned 2A1 B30.2 and 3A1 B30.2 complex, BTN2A1 B30.2 (20 µM) and BTN3A1 B30.2 (10 µM) were incubated in the buffer at a 2:1 ratio and loaded into the sample cell, and the injection syringe was filled with the buffer containing 100 µM HMBPP.

For other pAgs, their concentrations were adjusted to 500 µM, BTN2A1 B30.2 and BTN3A1 B30.2 were adjusted to 100 µM and 50 µM at a 2:1 ratio.

For the experiments testing the binding of BTN2A1/2A2 B30.2 domain (including their mutants) to the BTN3A1 B30.2 domain with or without HMBPP, BTN3A1 B30.2 (100 µM) and HMBPP (300 µM) were incubated in the buffer at a 1:3 ratio and loaded into the sample cell, and the injection syringe was filled with the buffer containing 1 mM BTN2A1/2A2 B30.2 (including their mutants).

For the BTN2A1 B30.2 domain binding to the BTN3A1 B30.2 domain in the presence of DMAPP or IPP, the concentration of BTN2A1 B30.2 was adjusted to 2 mM, the concentrations of BTN3A1 B30.2 and DMAPP (or IPP) were adjusted to 200 µM and 2000 µM at a 1:10 ratio.

### Size-exclusion chromatography multi-angle light scattering (SEC-MALS)

SEC-MALS experiments were carried out using a Wyatt Dawn Heleos II multiangle light scattering detector (Wyatt Technology) coupled to an AKTA Purifier UPC10 FPLC protein purification system and a Superdex 200 size exclusion column (GE Healthcare). Protein molecular weights of the individual peaks observed in the size-exclusion chromatograms were analyzed by static light scattering (SLS) in conjunction with their corresponding refractive indices, using an online refractometer connected downstream of the SLS detector (Wyatt Optilab rEX). 2A1 B30.2 (5 mg/mL) and 3A1 B30.2 (5 mg/mL) were prepared in a buffer (20 mM HEPES pH 7.5, 150 mM NaCl). HMBPP was prepared in 0.5 mM. The experiments were carried out with a running buffer (20 mM HEPES pH 7.5, 150 mM NaCl) at a flow rate of 0.5 mL/min. A standard value of the refractive index, dn/dc = 0.185 mL/g, was used for all proteins.

### Molecular dynamics simulation

The crystal structures of HMBPP and DMAPP-bound complexes were prepared using Protein Preparation Wizard (Schrödinger Release 2021-1) using default settings. The apo complex was built by removing the HMBPP ligand after the preparation. All three systems were immersed in an orthorhombic SPC water box with 10 Å of buffer width and neutralized by randomly placing sodium ions in the water box. The prepared simulation systems were relaxed and simulated at 300K and 1 atm using Desmond and OPLS4 force field (Schrödinger Release 2021-1). We simulated HMBPP and DMAPP bound to the 3A1-2A1 complex as well as the apo form of the 3A1-2A1 complex for 100 ns.

### Quantitative model for Vγ9Vδ2 T cell activation by phosphoantigens

For the thirteen library compounds, a quantitative model for their activities for sensitizing MIA PaCa-2 cells to Vγ9Vδ2 T cell killing (EC_50_) using a partial least-square method to regress binding affinities to the pre-conditioned 2A1-3A1 complexes (*K*_D_) and logP, against the cell pEC_50_ results. That is: pEC_50_ (predicted) = a·p*K*_D_ + b·logP + c, where pEC_50_ is -log_10_EC_50_, p*K*_D_ is -log_10_*K*_D_, and a-c are coefficients. EC_50_ values were determined using the cell killing assays; *K*_D_ values were determined by ITC and logP values were calculated using Chemdraw.

### Synthetic aspects

HMBPP (**1**), DMAPP (**2**), IPP (**3**) and HMBPP analogs were synthesized as described previously (*39-41*).

#### (*E*)-4-hydroxy-3-methylbut-2-en-1-yl diphosphate (1, HMBPP)

^1^H NMR (400 MHz, D_2_O) δ ppm 5.63 (t, *J* = 6.8 Hz, 1H), 4.50 (dd, *J_1_* = *J_2_* = 7.2 Hz, 2H), 3.99 (s, 2H), 1.68 (s, 3H); ^31^P NMR (162 MHz, D_2_O) δ ppm -6.56 (d, *J* = 20.0 Hz, 1P), -10.38 (d, *J* = 20.0 Hz, 1P). HRMS (m/z) [M-H]^+^ calculated 260.9923, found 260.9929.

#### 3-methylbut-2-en-1-yl diphosphate (2, DMAPP)

^1^H NMR (400 MHz, D_2_O) δ ppm 5.43 (t, *J* = 7.24 Hz, 1H), 4.44 (t, *J* = 6.88 Hz, 2H), 1.75 (s, 3H), 1.70 (s, 3H); ^31^P NMR (162 MHz, D_2_O) δppm -8.41 (d, *J* = 21.38 Hz, 1P), -10.55 (d, *J* = 21.38 Hz, 1P). HRMS (m/z) [M-H]^+^ calculated 244.9980, found 244.9978.

#### 3-methylbut-3-en-1-yl diphosphate (3, IPP)

^1^H NMR (400 MHz, D_2_O) δ ppm 4.82 (m, 2H), 4.04 (t, *J* = 6.70 Hz, 2H), 2.38 (t, *J* = 6.7 Hz, 2H), 1.75 (s, 3H); ^31^P NMR (162 MHz, D_2_O) δ ppm -9.49 (d, *J* = 20.90 Hz, 1P), -10.79 (d, *J* = 20.9 Hz, 1P). HRMS (m/z) [M-H]^+^ calculated 244.9980, found 244.9981.

#### (*E*)-4-hydroxy-3-ethyl-but-2-enyl diphosphate (4)

^1^H NMR (400 MHz, D_2_O) δ ppm 5.55 (t, *J* = 6.80 Hz, 1H), 4.56 (dd, *J_1_* = *J_2_* = 7.20 Hz, 2H), 4.01 (s, 2H), 2.07 (q, *J* = 7.60 Hz, 2H), 0.92 (t, *J* = 7.60 Hz,3H); ^31^P NMR (162 MHz, D_2_O) δ ppm -8.66 (d, *J* = 19.44 Hz, 1P), -10.57 (d, *J* = 19.44 Hz, 1P). HRMS (m/z) [M-H]^+^ calculated 275.0086, found 275.0086.

#### (*E*)-3-(hydroxymethyl)hexa-2,5-dienyl diphosphate (5)

^1^H NMR (400 MHz, D_2_O) δ ppm 5.91-5.87 (m, 1H), 5.64 (t, *J* = 6.80 Hz, 1H), 5.16-5.10 (m, 2H), 4.54 (dd, *J_1_* = *J_2_* = 6.80 Hz, 2H), 4.12 (s, 2H), 2.91 (d, *J* = 6.40 Hz, 2H); ^31^P NMR (162 MHz, D_2_O) δ ppm -9.17 (d, *J* = 21.1 Hz, 1P), -10.66 (d, *J* = 21.1 Hz, 1P). HRMS (m/z) [M-H]^+^ calculated 287.0081, found 287.0086.

#### (*E*)-3-(hydroxymethyl)undec-2-enyl diphosphate (6)

^1^H NMR (400 MHz, D_2_O) δ ppm 5.64 (t, *J* = 6.12 Hz, 1H), 4.55 (dd, *J_1_* = *J_2_* = 6.72 Hz, 2H), 4.06 (s, 2H), 2.13 (t, *J* = 7.32 Hz, 2H), 1.39 - 1.28 (m, 14H), 0.85 (t, *J* = 6.12 Hz,3H); ^31^P NMR (162 MHz, D_2_O) δ ppm -10.5 (d, *J* = 19.44 Hz, 1P), -10.8 (d, *J* = 19.44 Hz, 1P). HRMS (m/z) [M-H]^+^ calculated 359.1021, found 359.1025.

#### (*E*)-4-hydroxy-3-(4-methylbenzyl)but-2-enyl diphosphate (7)

^1^H NMR (400 MHz, D_2_O) δ ppm 7.09-7.07 (m, 4H), 5.75 (t, *J* = 6.80 Hz, 1H), 4.59 (dd, *J_1_* = *J_2_* = 6.40 Hz, 2H), 3.85 (s, 2H), 3.37 (s, 2H), 2.18 (s, 3H); ^31^P NMR (162 MHz, D_2_O) δ ppm -7.61 (d, *J* = 21.10 Hz, 1P), -10.44 (d, *J* = 21.10 Hz, 1P). HRMS (m/z) [M-H]^+^ calculated 351.0397, found 351.0399.

#### (*E*)-4-hydroxy-3-methylpent-2-en-1-yl diphosphate (8)

^1^H NMR (400 MHz, D_2_O) δ ppm 5.61 (t, *J* = 6.64 Hz, 1H), 4.48 (t, *J=*6.96 Hz, 2H), 4.23 (q, *J* = 6.60 Hz, 1H), 1.65 (s, 3H), 1.21 (d, *J* = 6.52 Hz, 3H); ^31^P NMR (162 MHz, D_2_O) δ ppm -8.07 (d, *J* = 21.06 Hz, 1P), -10.51 (d, *J* = 21.06 Hz, 1P). HRMS (m/z) [M-H]^+^ calculated 275.0086, found 275.0095.

#### (*E*)-((hydroxy((4-hydroxy-3-methylbut-2-en-1-yl)oxy)phosphoryl)methyl)phosphonic acid (9)

^1^H NMR (400 MHz, D_2_O) δ ppm 5.63 (t, *J* = 6.64 Hz, 1H), 4.49 (t, *J*= 7.36 Hz, 2H), 4.02 (s, 2H), 2.15 (t, *J*= 19.76 Hz, 2H), 1.71 (s, 3H). ^31^P NMR (162 MHz, D_2_O) δ ppm 18.37 (d, *J* = 9.56 Hz, 1P), 15.06 (d, *J* = 9.56 Hz, 1P). HRMS (m/z) [M-H]^+^ calculated 259.0137, found 259.0137.

#### (*E*)-(dichloro(hydroxy((4-hydroxy-3-methylbut-2-en-1yl)oxy)phosphoryl)methyl diphospha-te (10)

^1^H NMR (400 MHz, D_2_O) δ ppm 5.55 (t, *J* = 6.2 Hz, 1H), 4.60 (dd, *J*_1_= *J*_2_= 7.4 Hz, 2H), 3.90 (s, 2H), 1.59 (s, 3H). ^31^P NMR (162 MHz, D_2_O) δ ppm 10.92 (d, *J* = 9.0 Hz, 1P), 8.07 (d, *J* = 9.0 Hz, 1P). HRMS (m/z) [M-H]^+^ calculated 326.9357, found 326.9359.

#### (*E*)-(chloro(hydroxy((4-hydroxy-3-methylbut-2-en-1-yl)oxy)phosphoryl)methyl)phosphonic acid (11)

^1^H NMR (400 MHz, D_2_O) δ ppm 5.63 (t, *J* = 6.2 Hz, 1H), 4.56 (dd, *J*_1_ = *J*_2_ = 7.4 Hz, 2H), 4.0 (s, 2H), 3.8 (dd, *J*_1_ = *J*_2_ = 15.4 Hz, 1H), 1.70 (s, 3H). ^31^P NMR (162 MHz, D_2_O) δ ppm 13.99 (d, *J* = 9.0 Hz, 1P), 9.31 (d, *J* = 9.0 Hz, 1P). HRMS (m/z) [M-H]^+^ calculated 345.0667, found 345.0669.

#### (*E*)-(difluoro(hydroxy((4-hydroxy-3-methylbut-2-en-1-yl)oxy)phosphory)methyl)phosphoni -c acid (12)

^1^H NMR (400 MHz, D_2_O) δ ppm 5.61 (t, *J* = 6.4 Hz, 1H), 4.58 (dd, *J*_1_= *J*_2_= 7.28 Hz, 2H), 3.99 (s, 2H), 1.68 (s, 3H). ^31^P NMR (162 MHz, D_2_O) δ ppm 12.86 (m, 1P), 7.52 (m, 1P). HRMS (m/z) [M-H]^+^ calculated 294.9948, found 294.9952.

#### (*E*)-(fluoro(hydroxy((4-hydroxy-3-methylbut-2-en-1-yl)oxy)phosphoryl)methyl)phosphonic acid (13)

^1^H NMR (400 MHz, D_2_O) δ ppm 5.61 (t, *J* = 6.6 Hz, 1H), 4.64 (dd, *J*_1_= *J*_2_= 12.2 Hz, 1H), 4.54 (dd, *J*_1_= *J*_2_= 7.28 Hz, 2H), 3.98 (s, 2H), 1.68 (s, 3H). ^31^P NMR (162 MHz, D_2_O) δ ppm 12.86 (dd, *J*_1_=11.2 Hz, *J*_2_= 62.5 Hz, 1P), 7.52 (dd, *J*_1_= 11.2 Hz, *J*_2_= 58.0 Hz, 1P). HRMS (m/z) [M-H]^+^ calculated 277.0167, found 277.0169.

---

**Fig. S1.**
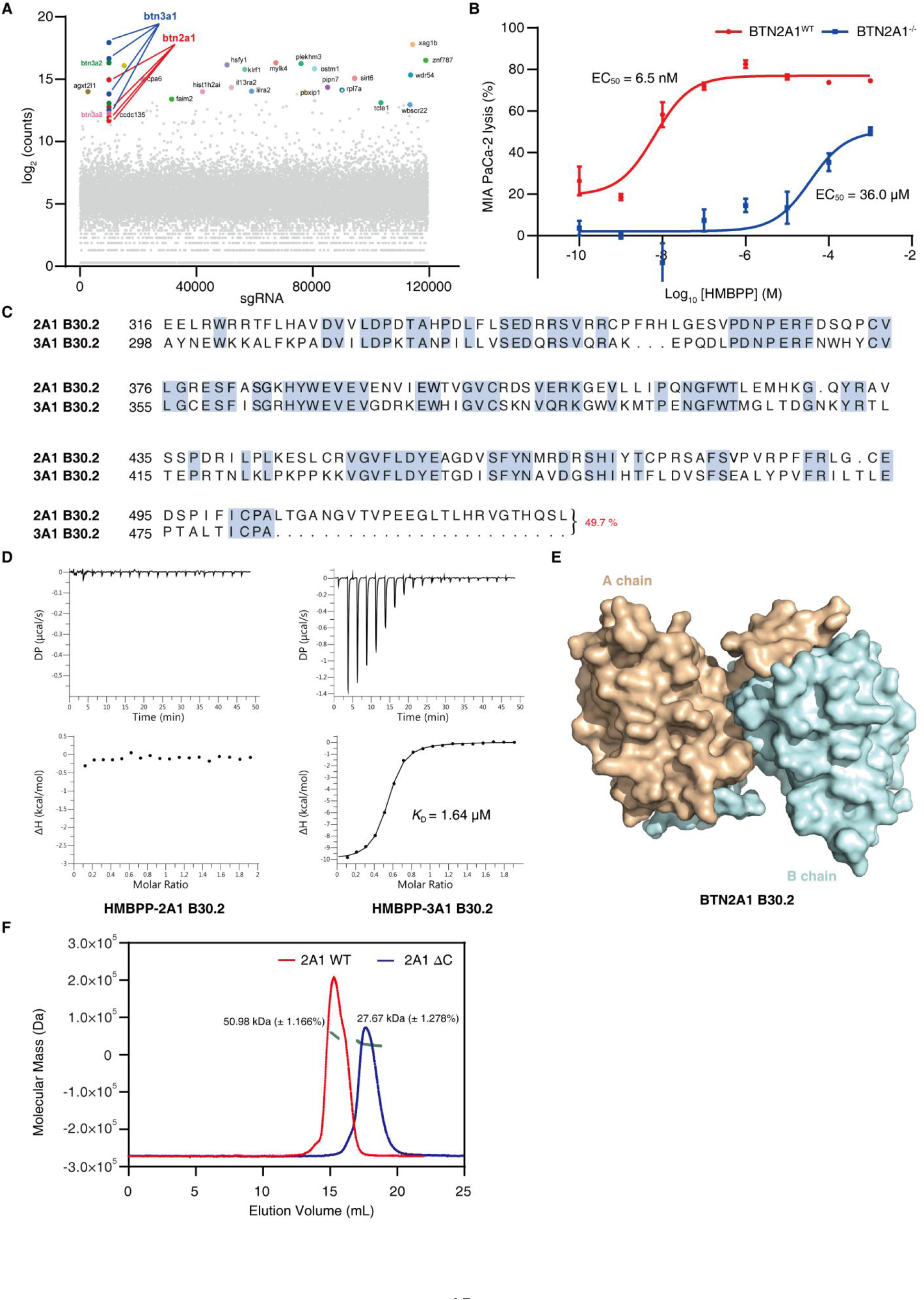
HMBPP binds to the BTN3A1 B30.2 domain but not to BTN2A1’s B30.2 domain. **(A)** MIA PaCa-2 cells transfected with a whole-genome sgRNA library were pretreated with HMBPP (10 nM) for 4 h, and then co-incubated with Vγ9Vδ2 T cells 7 consecutive times to enrich genes related to killing processes. BTN3A1 and BTN2A1 (sgRNA ≥ 4) were identified in the screen. **(B)** Cytotoxicity of Vγ9Vδ2 T cells towards BTN2A1^WT^ (red) or BTN2A1^-/-^ (blue) MIA PaCa-2 cells treated with HMBPP (100 pM to 1 mM). **(C)** Sequence alignment of the B30.2 domains of 2A1 and 3A1 (49.73% similarity). **(D)** ITC results for HMBPP binding to 2A1 B30.2 (left) and binding to 3A1 B30.2 (right). **(E)** The dimer interface of 2A1 B30.2 buried approximately 1923.6 Å^2^, (PDBePISA server: https://www.ebi.ac.uk/pdbe/pisa/), which accounts for 16% of the total surface area of the monomer. **(F)** SEC-MALS analysis of 2A1 B30.2 and 2A1 B30.2-ΔC. 2A1 B30.2 (in red) exists mainly as a dimer in solution. The 2A1 B30.2-ΔC domain (in blue) exists mainly as a monomer in solution.

**Fig. S2.**
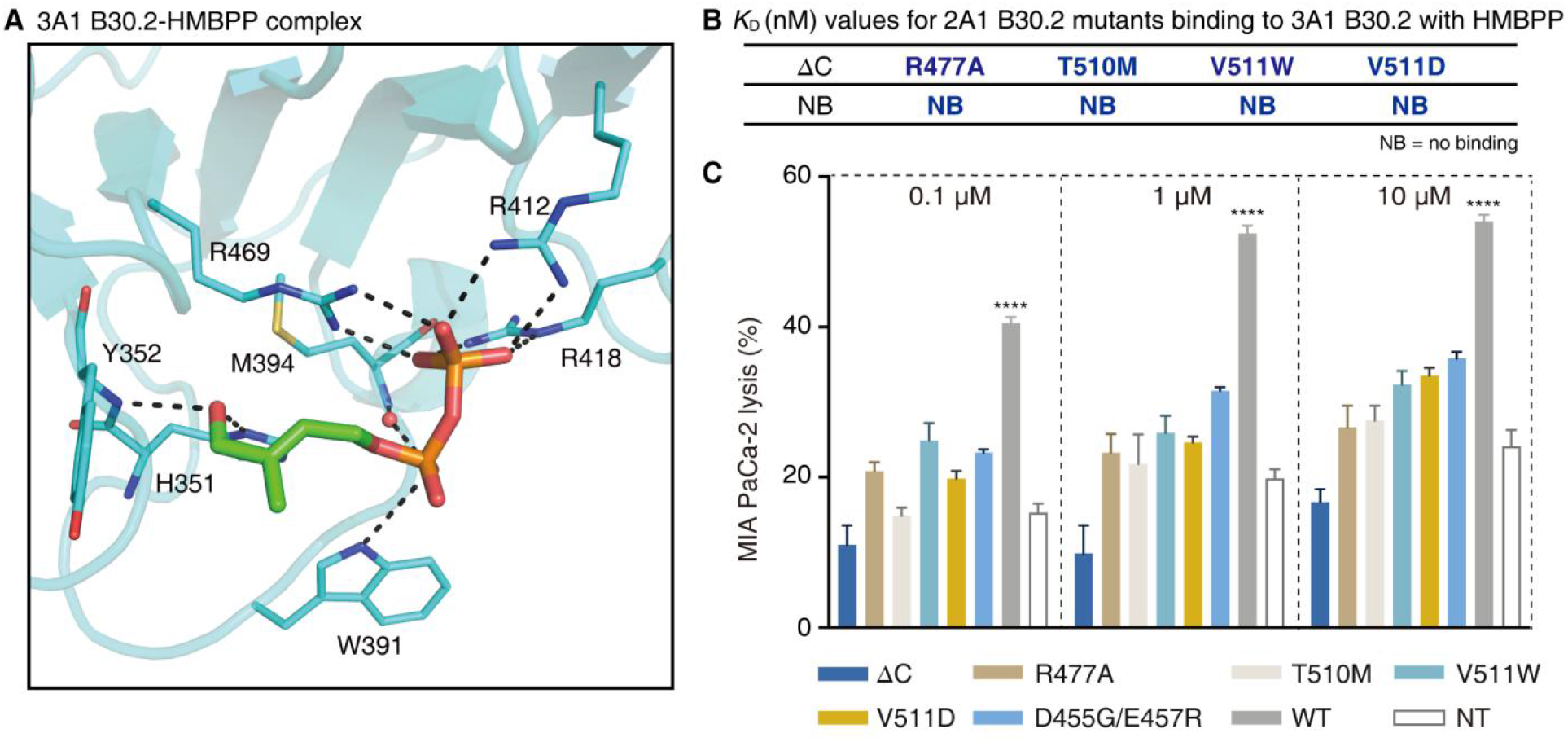
Mutagenesis of BTN2A1 residues reduces its intracellular association with BTN3A1 and impairs HMBPP-induced Vγ9Vδ2 T cell activation. **(A)** An expanded view of HMBPP binding to 3A1 B30.2 (in cyan) (PDB code: 5ZXK), showing the same interactions as observed in the 3A1 B30.2-HMBPP-2A1 B30.2 complex (Fig. 2C, iv). HMBPP and protein residues are shown as sticks and lines, respectively. **(B)** ITC results for the indicated 2A1 B30.2 mutants binding to 3A1 B30.2 in the presence of HMBPP. **(C)** Cytotoxicity of Vγ9Vδ2 T cells towards BTN2A^-/-^ MIA PaCa-2 (2A1/2A2 KO) transfected with plasmids for the indicated 2A1 mutant variants, treated with HMBPP (0.1 µM, 1 µM and 10 µM). Data were analyzed by using two-tailed unpaired *t* tests. *n* = 4, \*\*\*\**P* < 0.0001. All error bars denote the SEM.

**Fig. S3.**
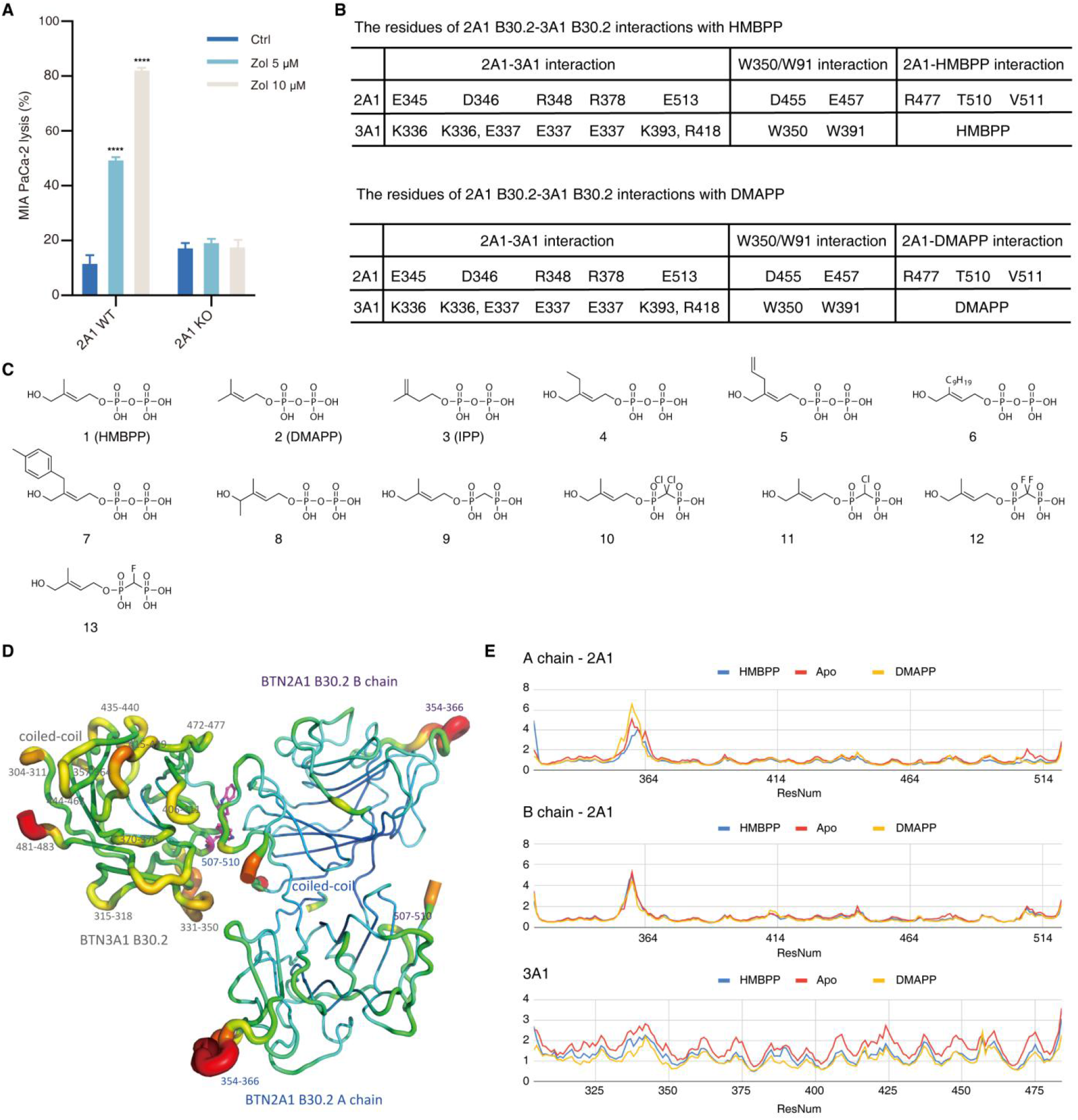
DMAPP and thirteen synthetic pAgs promote the 2A1-3A1 B30.2 association. **(A)** Cytotoxicity of Vγ9Vδ2 T cells towards BTN2A1^WT^ and BTN2A1^-/-^ MIA PaCa-2 cells, which were pretreated with zoledronate (Zol) for 24 h. Data were analyzed by using two-tailed unpaired *t* tests. *n* = 6, \*\*\*\**P* < 0.0001. All error bars denote the SEM. **(B)** Identification of residues from 2A1 B30.2 and 3A1 B30.2 which interact with HMBPP or DMAPP. **(C)** Chemical structures of natural and synthetic phosphoantigens (HMBPP analogs). **(D)** Apo-structure regions with RMSF > 1.5 Å in the molecular dynamics trajectory are mapped to the crystal structure (represented by 3A1 B30.2-HMBPP-2A1 B30.2 complex). Increasing thickness and red tube color represent regions with high RMSF values; the residue number of the segments are marked. Residues W350 and W391 are shown as sticks. **(E)** Cα-RMSF profiles for simulations with the apo-complex as well as for HMBPP-bound and DMAPP-bound complexes.

**Fig. S4.**
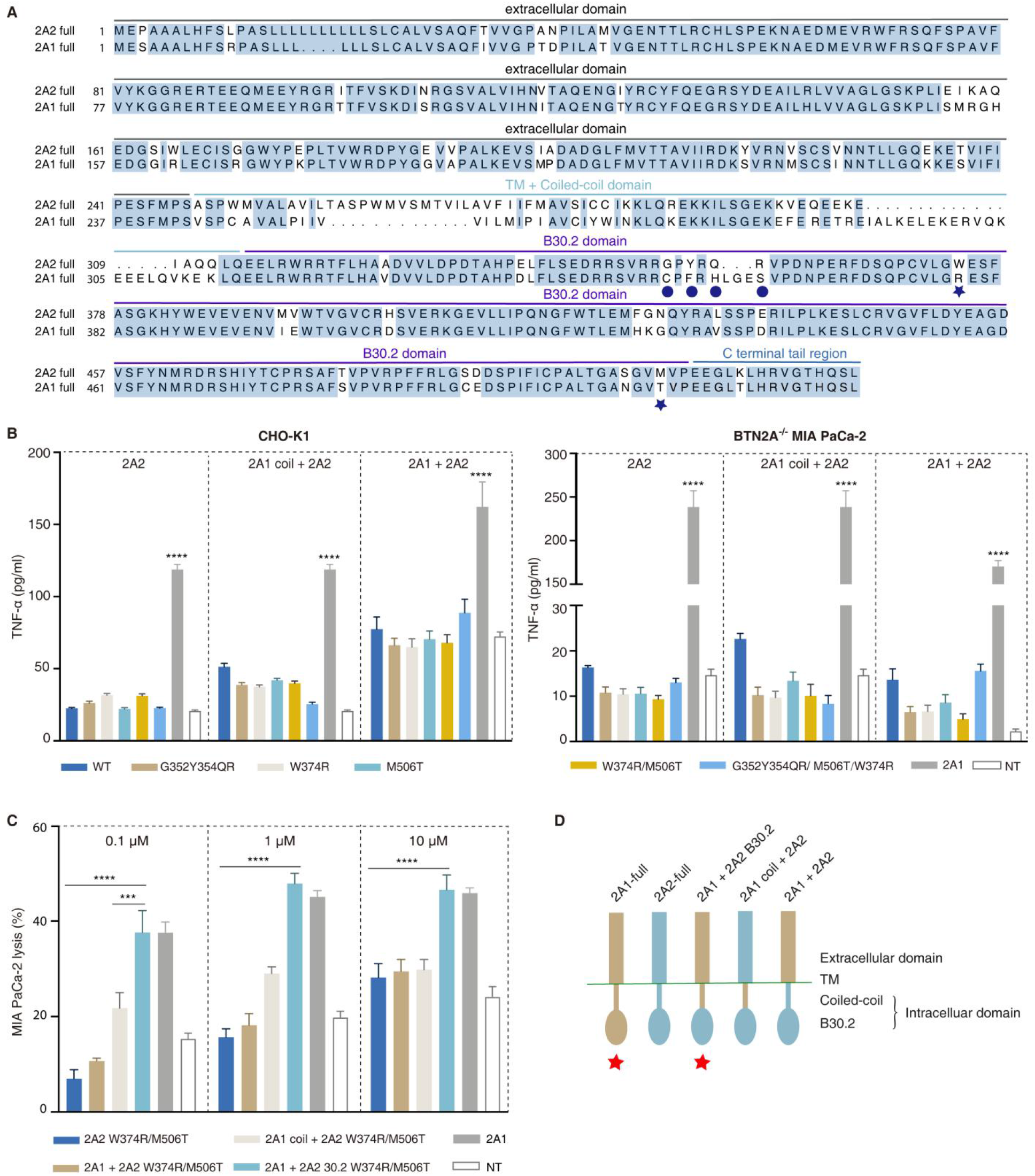
Mutation analysis of BTN2A2 and engineered variants for Vγ9Vδ2 T cell activation. **(A)** Sequence alignment of full-length 2A1 and 2A2 proteins. Sequence of BTN2A1 B30.2 and BTN2A2 B30.2 have 88.7% similarity. Blue stars indicate the residues in 2A2 (W374 and M506) analogous to 2A1 B30.2 residues that were identified to undergo functionally relevant interactions in the 2A1 B30.2-HMBPP-3A1 B30.2 complex structure. Note that 2A1 has an additional loop region (C353F355HLGES, in the blue circle) which is not present in the 2A2 structure (G352Y354QR). **(B)** TNF-α release by Vγ9Vδ2 T cells in response to zoledronate (10 µM) stimulation of 3A1^+^CD80^+^ CHO-K1 cells (left) or BTN2A^-/-^ MIA PaCa-2 cells (right) transfected with plasmids for the indicated combinations. Data were analyzed by using two- tailed unpaired *t* tests. *n* = 5, \*\*\*\**P* < 0.0001. All error bars denote the SEM. **(C)** Cytotoxicity of Vγ9Vδ2 T cells towards BTN2A^-/-^ MIA PaCa-2 cells transfected with plasmids for different combinations of 2A1 and 2A2 mutant variants, upon treatment with HMBPP (0.1 µM, 1 µM and 10 µM). Data were analyzed by using two-tailed unpaired *t* tests. *n* = 4, \*\*\**P* < 0.001 and \*\*\*\**P* < 0.0001. All error bars denote the SEM. **(D)** Sketches of mutations of different combinations of 2A1 and 2A2 (extracellular domain, the coiled-coil and TM, and B30.2 domain). The red star indicates that Vγ9Vδ2 T cells are activated.

**Table S1.** X-ray data collection and Refinement Statistics.

|  | BTN2A1 B30.2 | BTN3A1-BTN2A1-B30.2-HMBPP | BTN3A1-BTN2A1-B30.2-DMAPP | BTN2A2<br>B30.2<sup>W374R/M506T</sup> |
| --- | --- | --- | --- | --- |
| PDB code | 7EQ8 | 7EQQ | 7EQY | 7ER4 |
| Space group | P2<sub>1</sub>2<sub>1</sub>2 | P4<sub>1</sub> | P4<sub>1</sub> | P2<sub>1</sub>2<sub>1</sub>2<sub>1</sub> |
| Unit-cell |  |  |  |  |
| a (Å) | 73.957 | 89.823 | 90.153 | 55.673 |
| b (Å) | 78.889 | 89.823 | 90.153 | 59.843 |
| c (Å) | 41.682 | 169.335 | 167.918 | 127.612 |
| α / β / γ (°) | 90/90/90 | 90/90/90 | 90/90/90 | 90/90/90 |
| Resolution (Å) | 50-1.56(1.59-1.56) | 20-2.18(2.25-2.18) | 50-2.29(2.37-2.29) | 50-2.12(2.15-2.12) |
| Unique reflections | 35687(1647) | 69590(6594) | 60267(6050) | 24957(1012) |
| Redundancy | 12.70(10.10) | 13.44(10.72) | 13.70(13.40) | 9.62(5.48) |
| Completeness (%) | 99.6(93.5) | 99.4(94.2) | 100.0(100.0) | 99.9(100.0) |
| Average I/σ (I) | 27.89(3.5) | 21.9(4.9) | 25.67(4.00) | 11.6(2.45) |
| R<sub>merge</sub><sup>a</sup> (%) | 9.5(47.1) | 6.8(35.9) | 119.0(729.0) | 10.57(48.76) |
| No. of reflections | 33873(2378) | 65335(4567) | 57126(4179) | 23654(1697) |
| R<sub>work</sub> (95% of data) | 0.156(0.196) | 0.176(0.236) | 0.171(0.225) | 0.204(0.286) |
| R<sub>free</sub> (5% of data) | 0.173(0.211) | 0.213(0.302) | 0.216(0.272) | 0.260(0.303) |
| RMSD bonds [Å] | 0.014 | 0.012 | 0.011 | 0.008 |
| RMSD angles [°] | 1.806 | 1.757 | 1.710 | 1.545 |
| Dihedral angles |  |  |  |  |
| Most favored [%] | 97.5 | 97.2 | 96.4 | 97.4 |
| Allowed [%] | 2.5 | 2.7 | 3.6 | 2.6 |
| Disallowed [%] | 0.0 | 0.1 | 0.0 | 0.0 |
| No. of non-H atoms/ average B [Å<sup>2</sup>] |  |  |  |  |
| Protein chains | 1685/17.7 | 6132/46.6 | 6153/42.5 | 1602/33.3 |
| Water molecules | 249/29.8 | 382/52.7 | 343/47.1 | 134/35.8 |
| Ligand | 1/21.63 Zn<sup>2+</sup> | 30/33.98 HMBPP | 28/31.192 DMAPP |  |

Values in parentheses are for the highest resolution shell.
<sup>a</sup>R<sub>merge</sub> = $\sum_{\text{hkl}} \sum_i |I_i(\text{hkl}) - \langle I(\text{hkl}) \rangle| / \sum_{\text{hkl}} \sum_i I_i(\text{hkl})$ .

**Table S2.**
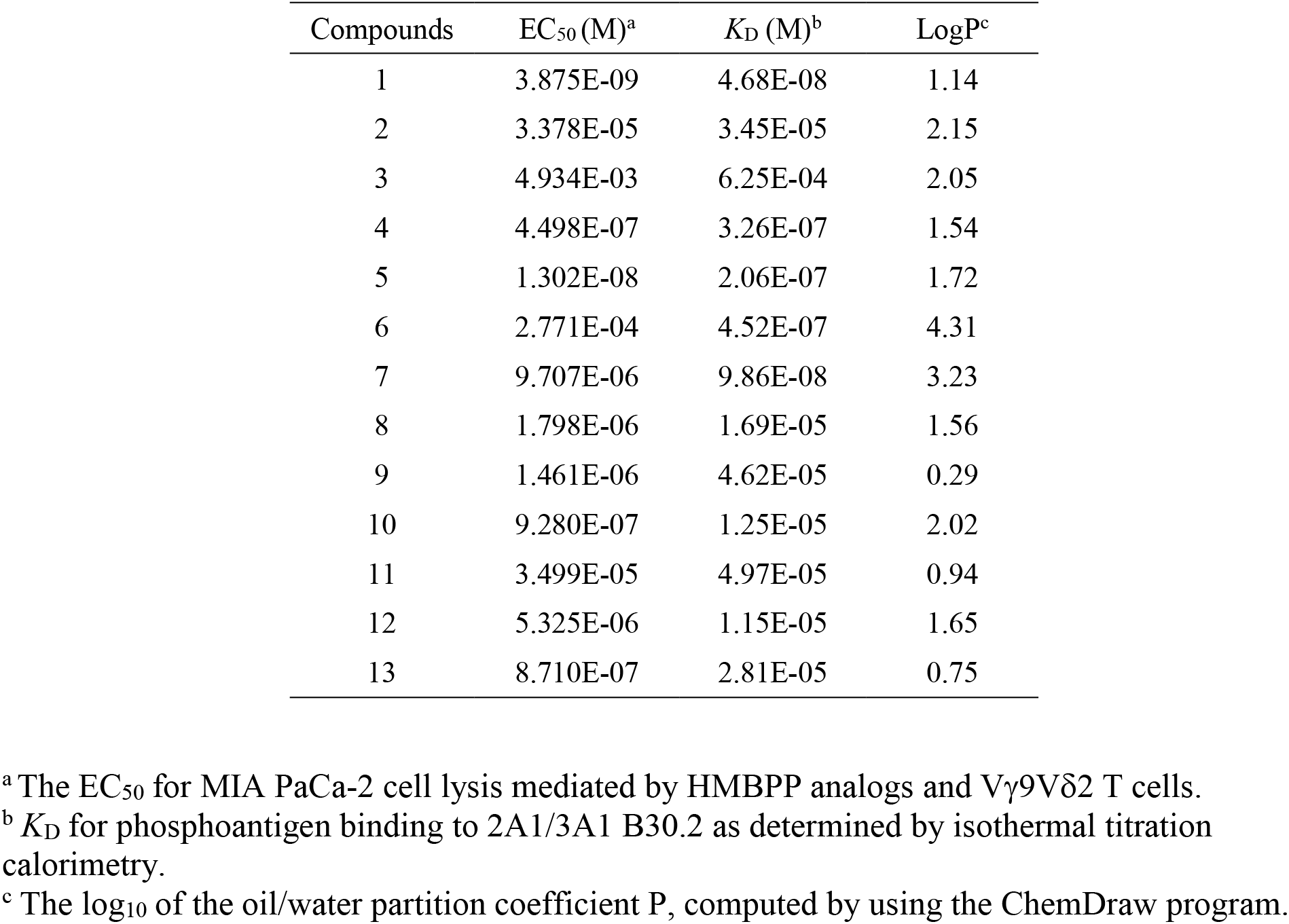
Summary of cell activity and binding affinity of HMBPP analogs.

| Compounds | EC<sub>50</sub> (M)<sup>a</sup> | K<sub>D</sub> (M)<sup>b</sup> | LogP<sup>c</sup> |
| --- | --- | --- | --- |
| 1 | 3.875E-09 | 4.68E-08 | 1.14 |
| 2 | 3.378E-05 | 3.45E-05 | 2.15 |
| 3 | 4.934E-03 | 6.25E-04 | 2.05 |
| 4 | 4.498E-07 | 3.26E-07 | 1.54 |
| 5 | 1.302E-08 | 2.06E-07 | 1.72 |
| 6 | 2.771E-04 | 4.52E-07 | 4.31 |
| 7 | 9.707E-06 | 9.86E-08 | 3.23 |
| 8 | 1.798E-06 | 1.69E-05 | 1.56 |
| 9 | 1.461E-06 | 4.62E-05 | 0.29 |
| 10 | 9.280E-07 | 1.25E-05 | 2.02 |
| 11 | 3.499E-05 | 4.97E-05 | 0.94 |
| 12 | 5.325E-06 | 1.15E-05 | 1.65 |
| 13 | 8.710E-07 | 2.81E-05 | 0.75 |

<sup>a</sup> The EC<sub>50</sub> for MIA PaCa-2 cell lysis mediated by HMBPP analogs and Vγ9Vδ2 T cells.
<sup>b</sup> K<sub>D</sub> for phosphoantigen binding to 2A1/3A1 B30.2 as determined by isothermal titration calorimetry.
<sup>c</sup> The log<sub>10</sub> of the oil/water partition coefficient P, computed by using the ChemDraw program.

**Table S3.**
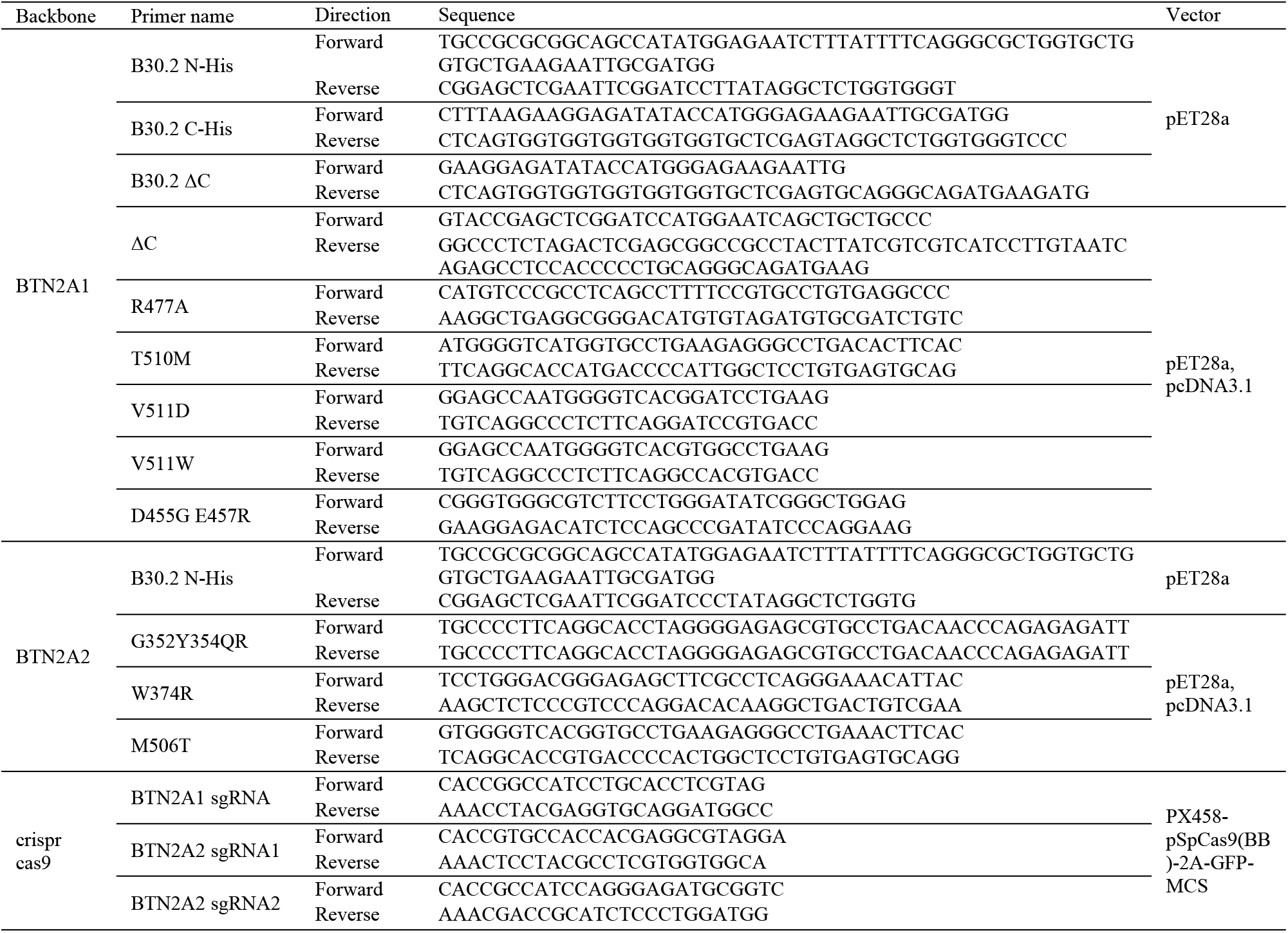
List of primers used.

